# Projection of orthogonal tiling from the retina to the visual cortex

**DOI:** 10.1101/2020.02.24.963785

**Authors:** Jaeson Jang, Min Song, Gwangsu Kim, Se-Bum Paik

## Abstract

In higher mammals, the primary visual cortex (V1) is organized into diverse tuning maps of visual features such as orientation, spatial frequency and ocular dominance. The topography of these maps is observed to intersect orthogonally, implying that a developmental principle for efficient tiling of sensory modules may exist. However, it remains unclear how such a systematic relationship among cortical tuning maps could develop. Here, we show that the orthogonal organization of tuning modules already exist in retinal ganglion cell (RGC) mosaics, and that this provides a blueprint of the orthogonal organization in V1. Firstly, from the analysis of multi-electrode recording data in V1, we found that the ON-OFF subregion distance of receptive fields varies periodically across the cortical surface, strongly correlated to ocular dominance and spatial frequency in the area. Further, the ON-OFF alignment angle, that is orthogonal to the ON-OFF distance, appears to correlate with orientation tuning. These suggest that the orthogonal organization in V1 may originate from the spatial organization of the ON-OFF receptive fields in the bottom-up projections, and this scenario was tested from analysis of the RGC mosaics data in monkeys and cats. We found that the ON-OFF RGC distance and ON-OFF angle of neighbouring RGCs are organized into a topographic tiling across mosaics, analogous to the orthogonal intersection of cortical tuning maps. These findings suggest that the regularly structured ON-OFF patterns mirrored from a retina may initiate efficient tiling of functional domains in V1.

**Highlights:**

- Orthogonal organization of visual tuning maps are observed in both V1 and the retina
- Cortical tuning maps are correlated with the profile of ON-OFF feedforward projections
- The profile of ON-OFF receptive fields varies periodically across the V1 surface
- Regularly structured RGC patterns initiate the orthogonal tiling of sensory modules in V1

---

The primary visual cortex (V1) of higher mammals is organized into a variety of functional maps of neural tuning such as preferred orientation, ocular dominance and spatial frequency. In the same cortical area, the gradient of orientation tuning intersects orthogonally with that of ocular dominance and preferred spatial frequency (1, 2) (Fig. 1A), resulting in an efficient tiling of diverse functional modules in V1 (2, 3) (Fig. 1B). Such structural correlation between the maps implies that there may exist a common principle of developing individual functional maps (2), but how such topographical relationships among different maps could arise in V1 remains unclear.

**Figure 1.**
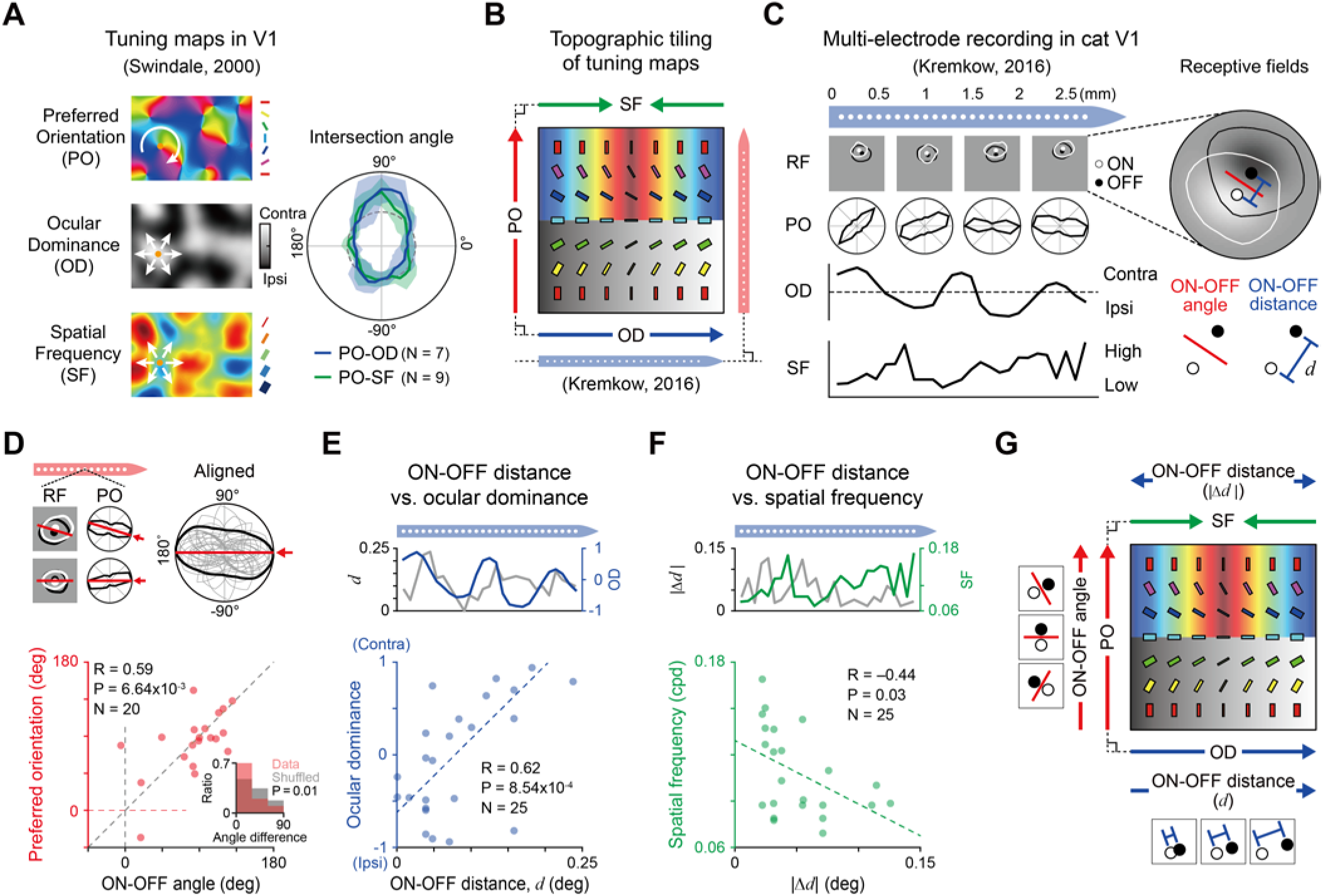
Topographic correlation between ON-OFF angle/distance and diverse cortical tunings. (A) Left: Diverse functional maps observed in V1. White solid arrows represent gradients in each map. Right: Orthogonal relationship between the gradients of diverse cortical tuning maps in (1–3)). N represents the number of map pairs examined (B) Efficient tiling of diverse function tunings illustrated. (C) Left: Receptive fields and functional tunings were recorded by the electrodes penetrating cat V1 (modified from Figure 2b of (5)). Right: ON-OFF angle and distance measured from the recorded receptive fields. (D) Top: ON-OFF angle estimated from local receptive field is correlated with the orientation tuning measured at each electrode with N = 20 data points. Bottom: Correlation between the ON-OFF angle and the preferred orientation. Inset: Angle difference between the two angles. The average difference is significantly smaller than that of the shuffled pairs (P = 0.01, N = 1,000 repeated trials). (E) Top: ON-OFF distance *(d)* and ocular dominance vary periodically across the cortical surface. Bottom: Correlation between *d* and ocular dominance. (F) Top: The deviation of ON-OFF distance from its average (|Δ*d*|) and the spatial frequency tuning measured by the same electrode. Bottom: Negative correlation between the preferred spatial frequency and |Δ*d*|. (G) In summary, diverse cortical tunings are constrained by the angle or the distance between ON and OFF feedforward afferents.

Important clues regarding the development of the maps were found in the studies of orientation tuning in feedforward afferents. It was reported that orientation tuning in a given cortical area is predictable from the angle alignment of the local ON and OFF thalamic afferents (4), and the change of orientation tuning across the cortical surface appears to correlate with the angle between ON and OFF receptive fields (5, 6). These observations support the idea of the statistical wiring model, in which the orientation tuning in V1 originates from the anisotropic profile in the local structure of ON and OFF retinal ganglion cell (RGC) mosaics (7–9), mirrored by feedforward mappings from a retina — via the lateral geniculate nucleus (LGN) (10) — to V1.

If this is so, how does the orthogonal organization between orientation tuning and other tunings arise in V1? Herein, we show that such an orthogonal relationship of tuning modules already exists in retinal mosaics, and that this can be mirrored to V1 to initiate the clustered topography of cortical organization. Our analysis of the published V1 recording data measured in cats (5) (Fig. 1C) show that ocular dominance and spatial frequency in V1 are correlated to the spatial separation of ON-OFF feedforward afferents (ON-OFF distance), intersecting orthogonally with the ON-OFF alignment angle (ON-OFF angle), which is correlated to orientation tuning. Interestingly, from analysis of the RGC mosaics data in cats (11, 12) and monkeys (13, 14), we found that the same type of orthogonal organization is observed in the retinal mosaics that provide bottom-up projections to V1. By combining observations in V1 recording and RGC mosaic data, we demonstrate that the bottom-up projection of regularly structured retinal afferents can solely initiate the efficient tiling of diverse tuning maps in the visual cortex.

## Correlation between ON-OFF afferents and cortical tunings

We analyzed recently published cat data (5), where the profile of V1 receptive fields and functional tuning were measured together across the cortical surface (Fig. 1C, left). In this dataset, measurements from two electrodes parallel or perpendicular to an ocular dominance column enabled an analysis of the relationship between the cortical receptive fields and the underlying cortical tuning of the preferred orientation and ocular dominance (red and blue electrodes in Fig. 1B). We found a systematic correlation between the spatial organization of local ON-OFF receptive fields and the ocular dominance/spatial frequency tuning at each location. From the observed receptive fields, we measured the angle and the distance between the center of ON and OFF subfields (Fig. 1C, right) and examined correlations with underlying functional tunings.

As reported in previous studies (4–6), we observed that orientation tuning measured at each electrode can be predicted by the angle between ON and OFF receptive fields (ON-OFF angle; Fig. 1D). More importantly, we found that the distance between ON and OFF centers (ON-OFF distance, *d*) periodically changes across the cortical surface (Fig. 1E, top). Notably, the spatial period of the ON-OFF distance variation was practically identical to that of the ocular dominance in the same cortical area (~2.9% difference; estimated by fitting to a sine curve). As a result, the observed ocular dominance appeared to correlate strongly to the ON-OFF distance (Fig. 1E, bottom; Pearson correlation coefficient, *N* = 25, *r* = +0.624, *P* = 8.54×10^-4^).

We also found that the ON-OFF distance and spatial frequency tuning are systematically correlated in the same data (Fig. 1F; Pearson correlation coefficient, *N* = 25, *r* = −0.46, *P* = 0.03). We found that the deviation of the ON-OFF distance from its average (|Δ*d*|; Fig. 1F, top, grey solid line) is correlated negatively with spatial frequency tuning estimated from the fast Fourier transform (FFT) analysis of V1 receptive fields (Fig. 1F, bottom). Considering the correlation between the ocular dominance and ON-OFF distance we observed (Fig. 1E, bottom), this topographic relationship is analogous to the previous observation that spatial frequency tuning and binocularity (1-|ocular dominance|) are correlated positively (2). These results suggest that the orthogonal organization of the spatial frequency and ocular dominance maps may be initialized by the spatial organization of the ON-OFF angle and distance in the bottom-up projections (Fig. 1G).

## Orthogonal organization of ON and OFF retinal ganglion cells

If the orthogonal tiling of tuning maps in a visual cortex originates from the spatial organization of ON-OFF afferents in feedforward projections (Fig. 1G), then it is expected that the ON-OFF angle and distance are already organized orthogonally in early stages of visual pathway, and that this spatial organization is mirrored to V1. Considering the precise retinotopic projection from the retina to V1, we hypothesized that the orderly arrangement of ON and OFF thalamic afferents to V1 originates from the spatial distribution of ON and OFF RGCs in retinal mosaics (Fig. 2A). To validate this scenario, we used RGC mosaics data previously measured in cats (11) (Fig. 2B, left) and examined whether the angle and distance between the local ON and OFF RGC mosaics are in orthogonal organization. Based on the statistical wiring model (7), we estimated the ON-OFF angle and distance of the sampled receptive field at each position of the RGC mosaics by assuming that RGCs are locally sampled to provide feedforward afferents in the local V1 area (Fig. 2B and 2C).

**Figure 2.**
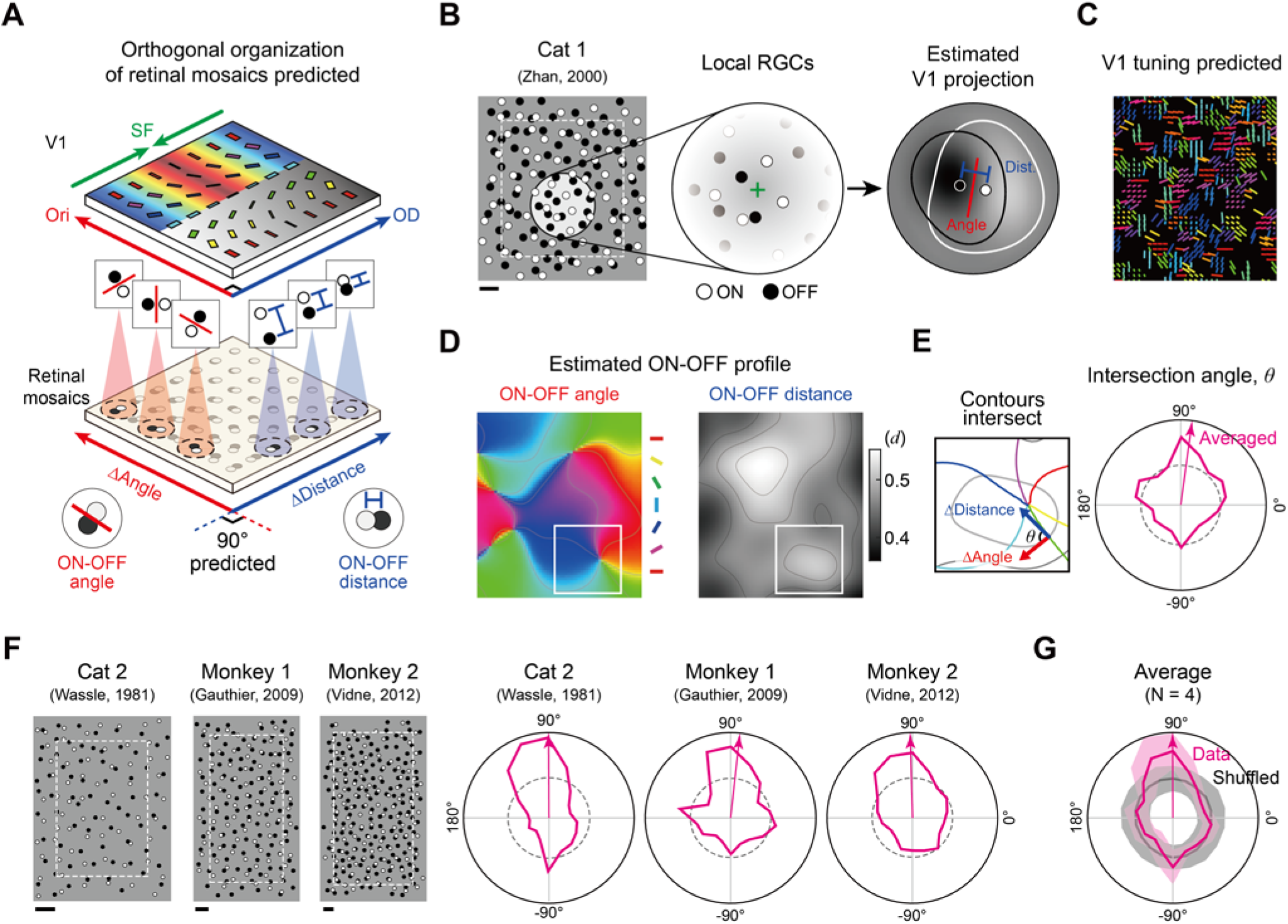
Orthogonal organization between ON-OFF angle and distance in RGC mosaics. (A) Because diverse cortical tunings change in the orthogonal direction and because each of them is topographically correlated with the ON-OFF angle or distance, the ON-OFF angle and distance in RGC mosaics are also predicted to change in the orthogonal direction. (B) Angle and distance between local ON and OFF receptive fields were estimated at each position of the measured RGC mosaic data. Boundary region (width = *1d_OFF_*, the expected average lattice distance of OFF RGC mosaics) was excluded for accurate estimation. The RGCs are statistically selected and summed with a two-dimensional Gaussian distribution (7). (C) Map of ON-OFF angle and distance measured within the white dashed square in (B). (D) Smoothed maps of ON-OFF angle and distance. (E) Left: Contours of ON-OFF angle (colourful solid lines) and distance (grey sold line) in the white solid square in smoothed maps. Right: Orthogonal intersection between gradients of ON-OFF angle and distance. Pink arrows indicate the average intersection angle. (F) Similar analyses for three other RGC mosaic data sets. (G) Averaged distribution of intersection angle shows a significant peak around 90°. Scale bar: 1*d_OFF_* measured in each mosaic.

We calculated the angle of the intersection between the ON-OFF angle and the distance in smoothed maps (Fig. 2D). As predicted by our scenario, we found that the angle shows a peak at approximately 90° (Fig. 2E). We repeated this analysis for four sets of RGC mosaic data of different species (n = 2 for cat (11, 12); n = 2 for monkey (13, 14)) and confirmed the orthogonal intersection between the ON-OFF angle and distance in all sets of tested data (Fig. 2F). The average distribution of all data sets showed a peak at 90° significantly higher than that of the control sets with the shuffled distribution of ON-OFF angle and distance (Fig. 2G; *P* = 0.01). These results show that the ON-OFF angle and distance in RGC mosaics intersect orthogonally across retinal space and this may provide the blueprint of orthogonal organization of functional tuning maps in V1.

## Discussion

Our results suggest that the orthogonal organization among the functional maps in V1 is initiated by regularly structured retinal afferents. Previously observed topographic correlation between the orientation pinwheels and ocular dominance (13) is also readily explained by our findings. Orientation pinwheels have been observed to be preferentially located around the center of monocular regions (16). From the observed correlation between the ocular dominance and the ON-OFF distance in our analysis (Fig. 1), together with our previous model approach in which orientation pinwheels emerge where the ON-OFF distance is locally minimized or maximized (15), both the orientation pinwheels and monocular regions are expected to be seeded by the same area in the RGC mosaics, resulting in the spatial overlap in the cortical space.

Overall, our results suggest that the regularly structured periphery circuits with simple feedforward wiring can provide a common developmental framework for diverse tuning circuitry in visual cortex.

## Materials and Methods

### Analysis of multi-electrode recordings

The multi-electrode data recorded from the cats were provided by Jose-Manuel Alonso ((5) and in personal communication) for the analysis in Figure 1. To remove high-frequency spatial noise, the receptive fields were filtered by a low-pass fermi filter in the frequency domain. The filter was designed according to

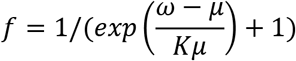

where *μ* is the threshold of the frequency (4 cpd) and *K* is the smoothness coefficient. The ON-OFF angle was estimated from the Fourier transform *Ψ(ω)* of the receptive fields. It is defined as *arg(μ)* /2, where

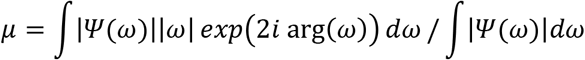

Samples for which either the ON or OFF subregion is entirely cancelled by the other component were excluded. The ON-OFF distance was defined as the distance between the center of the ON and OFF subregions. The location of the center of the ON and OFF subregions was defined as the strongest peak of each subregion. For multiple subregion samples, the largest subregion was chosen for analysis.

### Analysis of RGC mosaics

To estimate the ON-OFF angle and distance seeded at each location in the RGC mosaics, we modelled the cortical receptive fields by sampling the receptive fields of local ON and OFF RGCs. We assumed that the RGCs are statistically wired to cortical space with a two-dimensional (2D) Gaussian function with a standard deviation of *σ_con_*. The synaptic weighting between *i*^th^ RGC and *j*^th^ cortical sites, *w_ij_*, was defined as

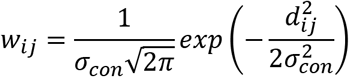

where *σ_con_* was set to 0.16–0.18 times the expected average lattice distance of OFF RGC mosaics (*d_OFF_*). The local receptive field was calculated at each vertex of a rectangular grid with a spacing distance of *0.1d_OFF_* in the RGC mosaics.

The receptive fields of RGCs and V1 neurons were defined as a center-surround model of 2D Gaussian and their linear sum, respectively. The standard deviation and the amplitude of the surrounding region was set to 3 and 0.55 times that of the center region (17).

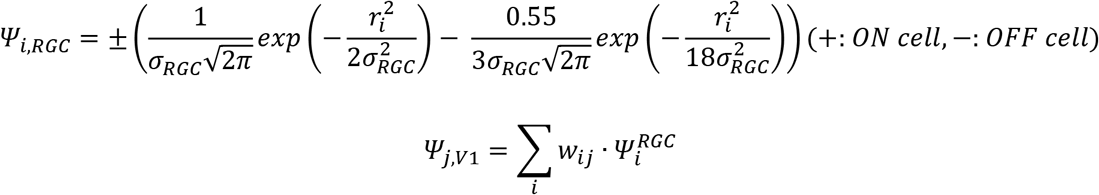

Here, *Ψ_i,RGC_* is the receptive field of *i^th^* RGC and *Ψ_j,v1_* is for the *j^th^* cortical site where *r_i_* is a distance vector from the center of *i^th^* RGC to each position of the visual field. The terms *σ_ON,RGC_* and *σ_OFF,RGC_* were set to 0.5 times the expected average lattice distance of ON and OFF RGC mosaics to satisfy the condition that the receptive fields of ON and OFF RGC mosaics cover all of the visual field (18). The ON-OFF angle and distance were measured between the center-of-mass of the modelled ON and OFF receptive fields and then the resultant maps were smoothed using a 2D Gaussian kernel with sigma of 0.8-1 *d_OFF_*.

## Acknowledgments

We are grateful to Jose-Manuel Alonso (State University of New York) for sharing receptive field data on the cat primary visual cortex. We also thank Dario Ringach, Jose-Manuel Alonso, Matteo Carandini, Daeyeol Lee and Andrea Benucci, for providing helpful comments in the early stages of this study. This work was supported by the National Research Foundation of Korea (NRF) grant funded by the Korea government (MSIT) (No. NRF-2019R1A2C4069863 and NRF-2019M3E5D2A01058328) (to S.P.).

## Author contributions

S.P. conceived the study. J.J., M.S. and S.P. designed the model. J.J. and M.S. performed the simulations. J.J., M.S., G.K. and S.P. analyzed the data. J.J., M.S. and S.P. wrote the manuscript. All authors discussed and commented on the manuscript.

